## Supplementary material for "Insect population dynamics under *Wolbachia*-induced cytoplasmic incompatibility: puzzle more than buzz in *Drosophila suzukii*": S1 Fig

Area 1	Area 2

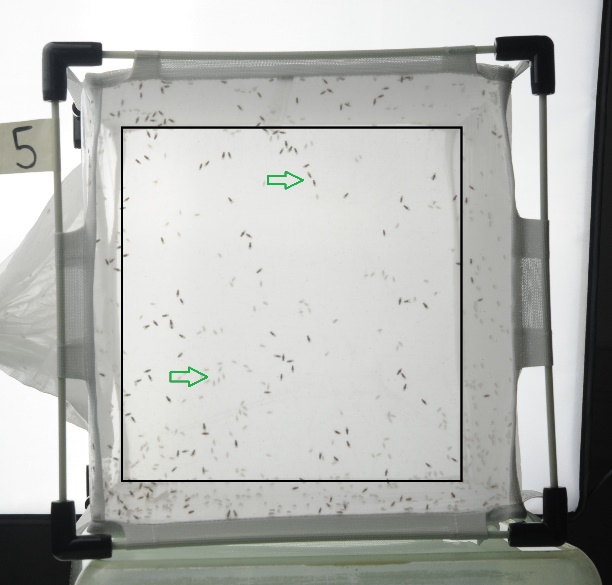

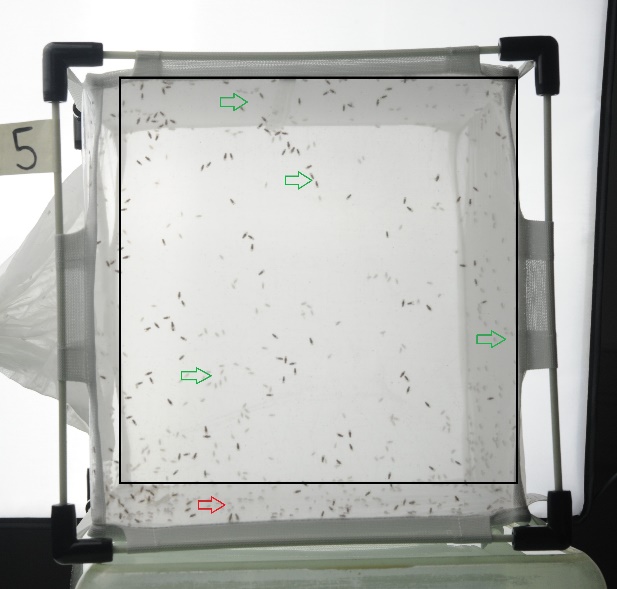

Area 3

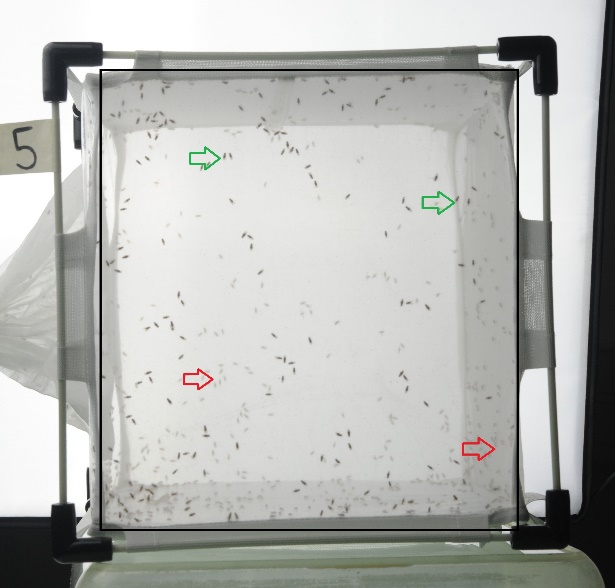

**Different areas used for the census method.** Black square represents area boundaries. Green and red arrows indicate which flies are counted and which are not. Area 1 : flies appearing both on the front and on the rear panels, Area 2 : individuals appearing on the whole surface except the left side and the bottom of the cage and Area 3 : only individuals appearing on the front panel.
