## Supplementary material for "Insect population dynamics under *Wolbachia*-induced cytoplasmic incompatibility: puzzle more than buzz in *Drosophila suzukii*": S2 Fig

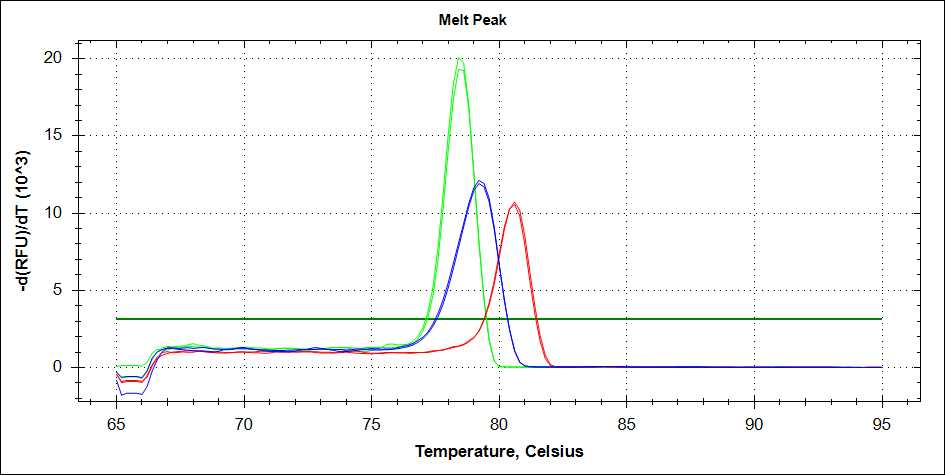

**Melting curves analysis of DNA fragments after PCR for different *Wolbachia* strains in *Drosophila suzukii*.**

Here, are plotted the negative first derivative of the fluorescence (-d(RFU)/dt (10 ^3^) *versus* temperature, peaks corresponding to the melting temperature Tm of *Wolbachia* DNA (Blue line: *w*Suz, green line: *w*Tei, red line: *w*Ha present in third transinfected line of *Drosophila suzukii*, which was not used for these experiments). FRU means relative fluorescence units.
