## Supplementary material for "Insect population dynamics under *Wolbachia*-induced cytoplasmic incompatibility: puzzle more than buzz in *Drosophila suzukii*": S3 Fig

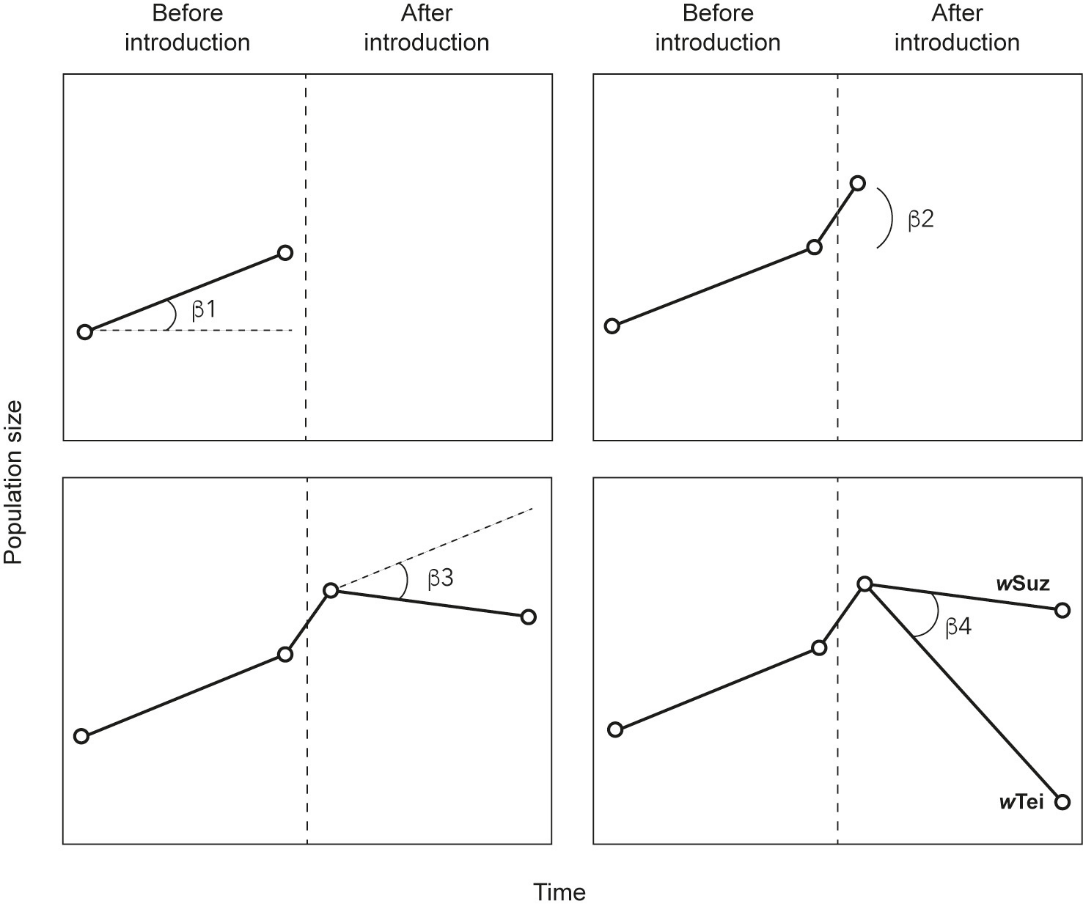

**Schematic representation of the statistical model fitted to time variations in number of *D. suzukii* in order to test the effect of introducing insects infected with either incompatible (*w*Tei) or compatible (*w*Suz) *Wolbachia* into a recipient population**.

Four hypotheses were tested, based on estimation of four model parameters. H1: parameter β_1_ population growth before introduction; populations were at carrying capacity and we therefore expected β_1_ = 0; H2: parameter β_2_ represents the immediate effect of introduction on population size; the number of flies introduced was set at 10% of the carrying capacity so we expected β_2_ > 0. H3: parameter β_3_ represents the effect of introduction on time variation of population size; if β_1_ = 0 and β_2_ > 0, we expected β_3_ < 0 reflecting a return to carrying capacity. H4: parameter β_4_ represents the effect of the *Wolbachia* strain introduced on the variation of population size after introduction; theoretical models predict that the introduction of an incompatible strain (*w*Tei) should result in a transient decrease in the population size, which should not be observed in control populations (introduction of *w*Suz), so that we expect β_4_ < 0.
