## Supplementary material for "Insect population dynamics under *Wolbachia*-induced cytoplasmic incompatibility: puzzle more than buzz in *Drosophila suzukii*": S1 Table

| ID | Fruit | Date | Nb_male_Suz | Nb_female_Suz | Nb_total_Suz |
| --- | --- | --- | --- | --- | --- |
| F1-02 | raspberry | 02/11/2021 | 0 | 0 | 0 |
| F5-22 | raspberry | 22/10/2021 | 0 | 0 | 0 |
| F5-02 | raspberry | 02/11/2021 | 0 | 0 | 0 |
| F4-02 | raspberry | 02/11/2021 | 0 | 0 | 0 |
| F10-18 | raspberry | 18/10/2021 | 0 | 0 | 0 |
| F3-29 | raspberry | 29/10/2021 | 0 | 0 | 0 |
| F2-22 | raspberry | 22/10/2021 | 0 | 0 | 0 |
| F6-22 | raspberry | 22/10/2021 | 0 | 0 | 0 |
| F10-27 | raspberry | 27/10/2021 | 0 | 0 | 0 |
| F2-27 | raspberry | 27/10/2021 | 0 | 0 | 0 |
| F2-04 | raspberry | 04/11/2021 | 1 | 0 | 1 |
| F1-29 | raspberry | 29/10/2021 | 0 | 1 | 1 |
| F2-29 | raspberry | 29/10/2021 | 1 | 0 | 1 |
| F8-18 | raspberry | 18/10/2021 | 0 | 1 | 1 |
| F4-04 | raspberry | 04/11/2021 | 1 | 0 | 1 |
| F3-02 | raspberry | 02/11/2021 | 0 | 1 | 1 |
| F9-02 | raspberry | 02/11/2021 | 0 | 1 | 1 |
| F8-22 | raspberry | 22/10/2021 | 1 | 0 | 1 |
| F11-18 | raspberry | 18/10/2021 | 0 | 1 | 1 |
| F7-22 | raspberry | 22/10/2021 | 1 | 0 | 1 |
| F12-18 | raspberry | 18/10/2021 | 0 | 2 | 2 |
| F4-22 | raspberry | 22/10/2021 | 1 | 1 | 2 |
| F7-27 | raspberry | 27/10/2021 | 2 | 0 | 2 |
| F9-22 | raspberry | 22/10/2021 | 0 | 2 | 2 |
| F5-18 | raspberry | 18/10/2021 | 1 | 2 | 3 |
| F8-02 | raspberry | 02/11/2021 | 0 | 3 | 3 |
| F8-27 | raspberry | 27/10/2021 | 3 | 0 | 3 |
| F7-18 | raspberry | 18/10/2021 | 2 | 1 | 3 |
| F13-18 | raspberry | 18/10/2021 | 2 | 2 | 4 |
| F14-18 | raspberry | 18/10/2021 | 1 | 3 | 4 |
| F3-27 | raspberry | 27/10/2021 | 1 | 3 | 4 |
| F4-27 | raspberry | 27/10/2021 | 3 | 1 | 4 |
| F6-27 | raspberry | 27/10/2021 | 3 | 1 | 4 |
| F3-22 | raspberry | 22/10/2021 | 0 | 4 | 4 |
| F1-22 | raspberry | 22/10/2021 | 4 | 1 | 5 |
| F5-27 | raspberry | 27/10/2021 | 2 | 3 | 5 |
| F9-27 | raspberry | 27/10/2021 | 5 | 1 | 6 |
| F3-04 | raspberry | 04/11/2021 | 4 | 3 | 7 |
| F1-04 | raspberry | 04/11/2021 | 4 | 4 | 8 |
| F3-18 | raspberry | 18/10/2021 | 5 | 3 | 8 |
| F7-02 | raspberry | 02/11/2021 | 3 | 5 | 8 |
| F2-18 | raspberry | 18/10/2021 | 7 | 2 | 9 |
| F1-18 | raspberry | 18/10/2021 | 6 | 5 | 11 |
| F6-18 | raspberry | 18/10/2021 | 7 | 5 | 12 |
| F9-18 | raspberry | 18/10/2021 | 7 | 6 | 13 |
| F1-27 | raspberry | 27/10/2021 | 10 | 5 | 15 |
| F4-18 | raspberry | 18/10/2021 | 10 | 9 | 19 |
| M7-18 | blackberry | 18/10/2021 | 0 | 1 | 1 |
| M4-18 | blackberry | 18/10/2021 | 0 | 1 | 1 |
| M2-22 | blackberry | 22/10/2021 | 1 | 0 | 1 |
| M5-04 | blackberry | 04/11/2021 | 1 | 0 | 1 |
| M3-20 | blackberry | 20/10/2021 | 3 | 2 | 5 |
| M8-22 | blackberry | 22/10/2021 | 3 | 2 | 5 |
| M8-04 | blackberry | 04/11/2021 | 1 | 4 | 5 |
| M3-16 | blackberry | 16/10/2021 | 1 | 4 | 5 |
| M1-20 | blackberry | 20/10/2021 | 4 | 2 | 6 |
| M4-04 | blackberry | 04/11/2021 | 3 | 3 | 6 |
| M7-04 | blackberry | 04/11/2021 | 2 | 4 | 6 |
| M5-22 | blackberry | 22/10/2021 | 4 | 3 | 7 |
| M3-18 | blackberry | 18/10/2021 | 3 | 5 | 8 |
| M1-18 | blackberry | 18/10/2021 | 5 | 7 | 12 |
| M5-16 | blackberry | 16/10/2021 | 8 | 9 | 17 |
| M4-22 | blackberry | 22/10/2021 | 9 | 11 | 20 |
| M4-16 | blackberry | 16/10/2021 | 9 | 12 | 21 |

**Number of adults *D. suzukii* emerging from fruits exposed in the field.**

Number of males and females *D. suzukii* emerged from raspberries and blackberries exposed in the field for 48h.
